# Reference-free identification and pangenome analysis of accessory chromosomes in a major fungal plant pathogen

**DOI:** 10.1101/2024.12.12.627383

**Authors:** Anouk C. van Westerhoven, Like Fokkens, Kyran Wissink, Gert Kema, Martijn Rep, Michael F. Seidl

## Abstract

Accessory chromosomes, found in some but not all individuals of a species, play an important role in pathogenicity and host specificity in fungal plant pathogens. However, their variability complicates reference-based analysis, especially when chromosomes are missing from reference genomes. Pangenome variation graphs offer a reference-free alternative for studying these chromosomes. Here, we constructed a pangenome variation graph for *Fusarium oxysporum*, a major fungal plant pathogen with a compartmentalized genome. To study accessory chromosomes, we constructed a chromosome similarity network and identified eleven conserved core chromosomes and many highly variable accessory chromosomes. Some of these are host-specific and are likely involved in determining host range, which we corroborate by analyzing nearly 600 *F. oxysporum* assemblies. By a reconstruction of pangenome variation graph per homologous chromosomes, we show that these evolve due to extensive structural variation as well as the exchange of genetic material between accessory chromosomes giving rise to these mosaic accessory chromosomes. Furthermore, we show that accessory chromosomes are horizontally transferred in natural populations. We demonstrate that pangenome variation graphs are a powerful approach to elucidate the evolutionary dynamics of accessory chromosomes in *F. oxysporum* and provides a computational framework for similar analyses in other species that encode accessory chromosomes.

## Introduction

The number of chromosomes of individuals in a single species is generally conserved. However, in various plants, animals, oomycetes, and fungi, a variable number of chromosomes is identified within a species. (D’Ambrosio *et al*., 2017). These accessory chromosomes often encode very few genes, with low transcriptional activity, and little effect on the phenotype (D’Ambrosio *et al*., 2017; Ma *et al*., 2017). In contrast, accessory chromosomes in various fungal plant pathogens encode genes with important roles in host infection (Ma *et al*., 2010; Habig *et al*., 2017; Torres *et al*., 2020; van Westerhoven *et al*., 2024a; Wei *et al*., 2024). These accessory chromosomes are often highly variable across individuals and show extensive presence / absence variation (Fouché *et al*., 2018; Stalder *et al*., 2022; van Westerhoven *et al*., 2024a). The presence of pathogenicity genes on these variable accessory chromosomes separates pathogenicity genes from house-keeping genes and has been thought to facilitate rapid adaptation to changing environments as well as the host immune system (Croll & McDonald, 2012; Torres *et al*., 2020).

The increasing availability of continuous, chromosome-level genome assemblies enabled the identification of accessory chromosomes in numerous fungi (Coleman *et al*., 2009; Haas *et al*., 2009; Ma *et al*., 2010; Rouxel *et al*., 2011; Goodwin *et al*., 2011; Dong *et al*., 2015; Henry *et al*., 2021; Wacker *et al*., 2023; van Westerhoven *et al*., 2024a; Barragan *et al*., 2024), and allows for comparative genomic approaches that help to elucidate the evolution of accessory chromosomes (Plissonneau *et al*., 2016; Henry *et al*., 2021; van Westerhoven *et al*., 2024a; Barragan *et al*., 2024). However, the analysis of accessory chromosomes is limited by reference-based analyses, where the variation of accessory chromosomes can only be determined by comparisons to a single reference genome. Because of the variability of accessory regions, the reference genome often lacks specific accessory regions that will not be analyzed. To capture the complete variation in a collection of genome sequences, including the accessory chromosomes, reference-free analysis strategies are needed. Recent advances in pangenome analysis methods can now be used to capture all genetic variation in a species (Tettelin *et al*., 2005; The Computational Pan-Genomics Consortium, 2018). One approach is the pangenome variation graph (Eizenga *et al*., 2020; Hickey *et al*., 2023), where nodes represent sequences and edges connect adjacent nodes; each path through the graph represents a single genome. This graph can be used as a framework to call variants in a population, annotate genes, and analyze chromosome structure (Hickey *et al*., 2020; Guarracino *et al*., 2023; Skiadas *et al*., 2024).

Thus far, pangenome variation graphs have been mainly constructed to analyze the human genome as well as some plant genomes (Gao *et al*., 2019; Wang *et al*., 2022; Guarracino *et al*., 2023). Only recently a pangenome variation graph for the oomycete plant pathogen *Peronospora effusa* has been constructed to study processes driving genome evolution (Skiadas *et al*., 2024). In this study the construction of the pangenome graph was guided by a reference genome (Hickey *et al*., 2020, 2023), which can offer important insights in organisms where the chromosome structure is largely conserved. However, pangenome variation graphs are more challenging to construct for fungal genomes that typically contain more chromosomal variations, especially in accessory chromosomes (Croll & McDonald, 2012; Fouché *et al*., 2018; Dijkstra *et al*., 2024), and fungal pangenome variation graphs have not been published. Due to the lack of co-linearity, the analysis of accessory chromosomes using pangenome variation graphs is not straightforward. Grouping the chromosomes into homologous groups (Garrison *et al*., 2023) prior to construction of a variation graph is a promising strategy to analyze chromosomes in dynamic fungal species. However, its unknown how homologous groups should be constructed for species carrying accessory chromosomes.

The fungal plant pathogen *Fusarium oxysporum* can infect various important crops (van Dam *et al*., 2016; Edel-Hermann & Lecomte, 2019; Armer *et al*., 2024) and is known to have a compartmentalized genome where accessory regions can span entire chromosomes or can be attached to conserved core chromosomes (Ma *et al*., 2010; van Dam *et al*., 2017; Armitage *et al*., 2018; Li *et al*., 2020; Henry *et al*., 2021; van Westerhoven *et al*., 2024a; Dijkstra *et al*., 2024). The core chromosomes are largely co- linear with limited large-scale variation such as deletions, insertions, inversions, or translocations (Ma *et al*., 2010; Fayyaz *et al*., 2023; van Westerhoven *et al*., 2024a). In contrast, accessory chromosomes are diverse even between strains infecting the same host (Henry *et al*., 2021; van Westerhoven *et al*., 2024a). Importantly, horizontal transfer of accessory chromosomes between strains can transfer pathogenicity between strains in the lab (Ma *et al*., 2010; van Dam *et al*., 2017; Li *et al*., 2020). Thus far, the genome structure and diversity of accessory chromosomes in *Fusarium oxysporum* has mainly been analyzed for strains infecting a specific host in comparison to the widely studied reference genome *F. oxysporum* f.sp*. lycopersici* strain 4287 *(Fol*4287) infecting tomato, which is known to have five accessory chromosomes (Ma *et al*., 2010). Little is known about occurrence, the frequency, and the variation of accessory chromosomes across the *Fusarium oxysporum* species complex.

The presence of a compartmentalized genome together with the availability of various continuous genome assemblies make *F. oxysporum* a suitable model to analyze the dynamics of accessory chromosomes using a pangenome variation graph. Here, we constructed a pangenome variation graph to analyze the evolution of accessory chromosomes in *F. oxysporum.* Some of these accessory chromosomes were specific to strains infecting the same host, suggesting that these play a role in determining host specificity. Furthermore, accessory chromosomes were composed of mixed stretches of accessory regions and, likely, evolved through recombination of accessory chromosomes. These findings offer insights into the evolution of accessory chromosomes and show how pangenome variation graphs can be applied to analyze chromosome dynamics.

## Results

### *Fusarium oxysporum* contains eleven core chromosomes and various accessory chromosomes

To study the diversity of core and accessory chromosomes in a large collection of diverse *F. oxysporum* strains, we obtained 73 highly continuous whole-genome assemblies (contig N50 > 2 Mb, <100 contigs) (Table S1). Together, these strains contain 1,261 contigs. Most (1,131 out of 1,261) contigs encode telomeric repeats on at least one end, and 446 represent complete chromosomes containing telomeric repeats on both sites, we therefore refer to all contigs as chromosomes in this manuscript. The dataset contains *F. oxysporum* strains assigned to 19 different formae speciales based on their capacity to infect specific host plants, three endophytes, and eight strains that cause plant disease but have not been assigned to a formae speciales (Fig. 1a,b). The strains have an average nucleotide similarity of 97.7%, and span all three known phylogenetic clades within the *F. oxysporum* species complex (Fig. 1a; O’Donnell *et al*., 1998; Maryani *et al*., 2019).

**Figure 1.**
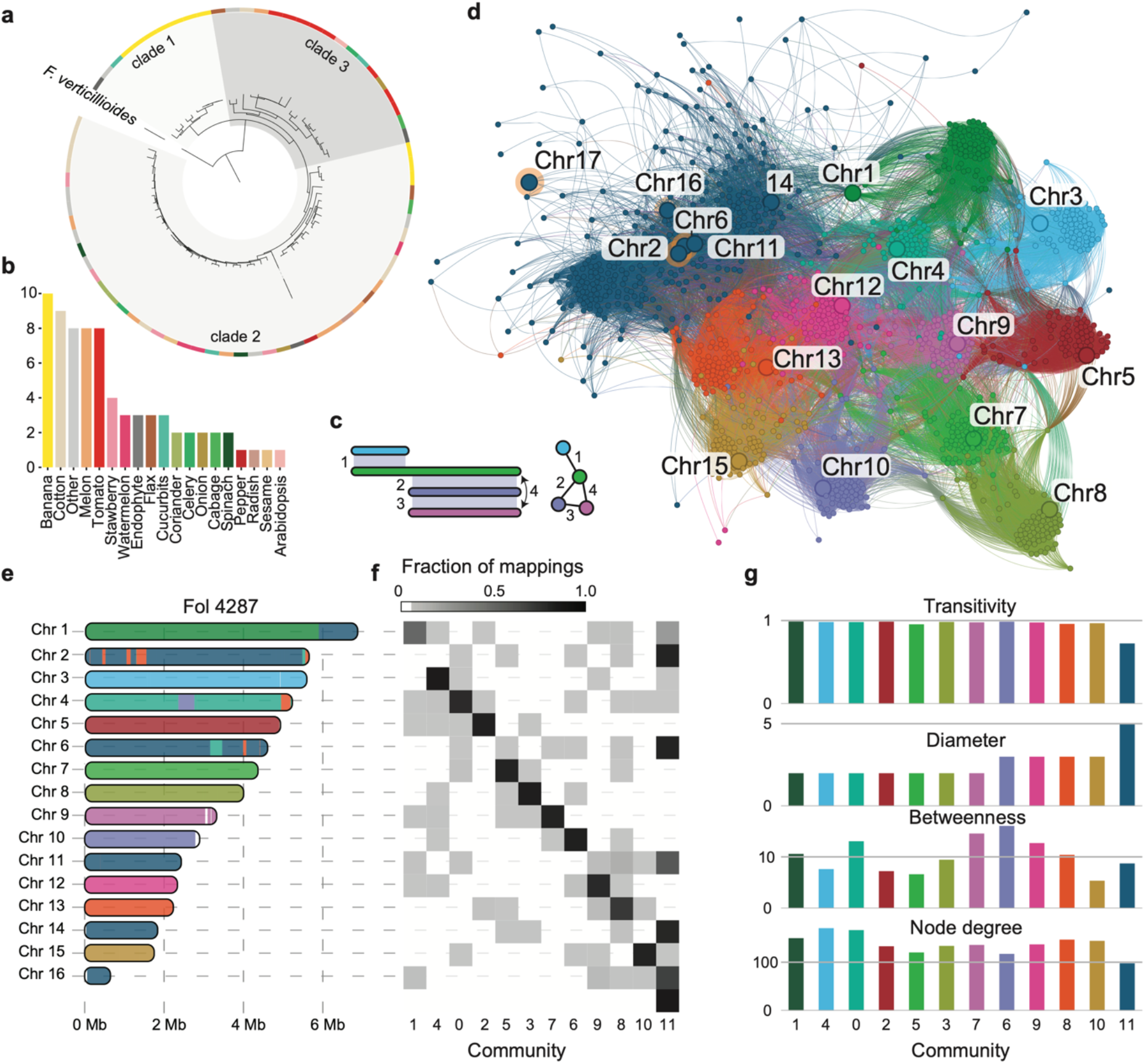
– Eleven core-chromosomes as well as a high number of accessory chromosomes are present in *Fusarium oxysporum*. **a**) Phylogenetic tree of 73 *Fusarium oxysporum* strains, based on 4,443 single-copy BUSCO genes. Different grey boxes highlight strains belonging to the three taxonomic lineages in the *F. oxysporum* species complex. The outer ring shows the different hosts that can be infected by a strain, colored according to (b). **b**) Distribution of plant host that can be infected by the 73 *F. oxysporum* strains. Endophytic strains as well as plant pathogens not assigned to a formae speciales (Other) are separated. **c**) The cartoon visualizes how sequences mappings between chromosomes are translated to a chromosome network. **d**) Chromosome network displays all pair-wise mappings between the chromosomes of the 73 *F. oxysporum* strains. Nodes represent chromosomes and edges represent mappings between chromosomes. Colors indicate the different communities that are detected. Chromosome names correspond to the chromosomes found in the *Fol*4287 reference genome, and these chromosomes are depicted by a large node in the graph. Orange circles highlight accessory chromosomes. All accessory chromosomes found in reference genome *Fol*4287 occur in cluster 11 (dark blue cluster). **e**) Mapping of communities to homologous chromosome in reference *Fol*4287, colored according to (d) **f**) Communities can be associated to specific chromosomes in *Fol4*287. The squares display the fraction of mappings of the *Fol*4287 chromosome (y-axis) with the corresponding community (x-axis). Core chromosomes map largely to a single community, while all accessory chromosomes are assigned to community 11. **g**) Network statistics of the different communities in the chromosome network. Colors correspond to the communities in the network, see (d).

To study the variation of chromosome content in these different *F. oxysporum* strains and to facilitate pangenome analysis, we first sought to detect groups of co-linear chromosomes based on all-vs-all homology mapping between all 1,261 chromosomes (Fig. 1c). We tried various thresholds for mapping, i.e., the percent identity and the length, to determine the optimal mapping strategy; the mappings need to be specific enough to map homologous chromosomes and to prevent spurious matches, yet sensitive enough to capture similarity between more distantly related chromosomes. These mappings between chromosomes were then translated into a chromosome network, and we used network properties to detect communities within the network that represent groups of homologous chromosomes (Fig. 1d). Since core chromosomes are expected to be conserved and co-linear, we expected to find communities of core chromosomes present in all strains, as well as several accessory chromosomes with a variable presence and absence pattern.

To analyze the distribution of the chromosomes over the communities, we determined the presence of chromosomes of the *F. oxysporum* f. sp. *lycopersici* reference genome *Fol*4287 in the communities (Fig. 1e-f). The core-chromosomes of *Fol*4287 are grouped into eleven separate communities, whereas the five accessory chromosomes (chromosomes 3,6,14,16, and 17) all group together into a single large community, which contained 342 chromosomes in total. Interestingly, two out of 73 strains do not encode any accessory chromosome (36102; infecting banana and Fo5; endophytic, table S2). The core-chromosomes on the other hand are all present in all 73 strains. Of the 11 core-chromosome communities, six communities contain a chromosome of all 73 strains and five communities miss one or two strains (Table S2). These missing chromosomes are the result of chromosomal rearrangements, either due to a mis-assembly or due to interchromosomal translocation. These fused chromosomes have similarity to multiple communities but are only assigned to the community with most matches. This also means that none of the strains lacks one of the core chromosomes (Table S2). Irrespective of the mapping threshold that determines the minimal length of mappings between chromosomes, we identified eleven core chromosomes, with one copy in all strains, as well as one community that contained all accessory chromosomes (Fig. 1d).

### Pangenome graph captures variation of *Fusarium oxysporum* and highlights differences between core and accessory chromosomes

All accessory chromosomes cluster in a single community (community 11; Fig. 1d), suggesting that many accessory chromosomes in *F. oxysporum* share genetic material. However, community 11 has a lower average node degree (89) and a larger diameter (5) compared with the core communities (average node degree 136; average diameter 2.2). Moreover, the nodes in this cluster are generally less well connected (transitivity 0.74 vs avg. transitivity of 0.97, figure 1g), collectively suggesting that some of the chromosomes in the community have limited similarity to each other but are connected through similarity with other chromosomes.

To further analyze the chromosome structure and the similarities between chromosomes, we constructed a pangenome variation graph using the PanGenome Graph Builder (PGGB). This graph consists of eleven subgraphs, one graph per community, and contains 500 Mb of genetic material split over 33,651,077 nodes connected through 48,127,154 edges. 2,559,224 nodes (18.6 Mb) are core (present in all 73 genomes), 13,404,553 nodes (41.1 Mb) are accessory, and 17,687,283 nodes (440.6 Mb) are unique to a single genome (Fig. 2a). The pangenome graph constructed for these 73 *F. oxysporum* genomes is open (Heaps Alpha = 0.327, Fig. 2b), meaning that every addition of a newly sequenced genome is expected to add new genetic material to the graph. Core, accessory, and unique material is found in different quantities in the different communities. The eleven core-chromosome communities contain genetic material present in all genomes. However, in five of these core-chromosome communities more accessory material is present (Fig. 2c), including communities containing the core-chromosomes 12, 13 and 15 of *Fol*4287. Interestingly, these chromosomes have been described as more variable than the other core chromosomes and have been previously referred to as ‘fast-core’ chromosomes (Fokkens et al. 2018). However, a similar pattern is also observed in chromosome 7 and 10, which are not considered part of the fast-core chromosomes, suggesting that these chromosomes similarly contain high amount of genomic variation. The amount of unique material is consistently low in the core chromosomes, and higher in the accessory community. As may be expected, the accessory community consists solely of accessory material (Fig. 2c), and not a single node is present in all accessory chromosomes.

**Figure 2.**
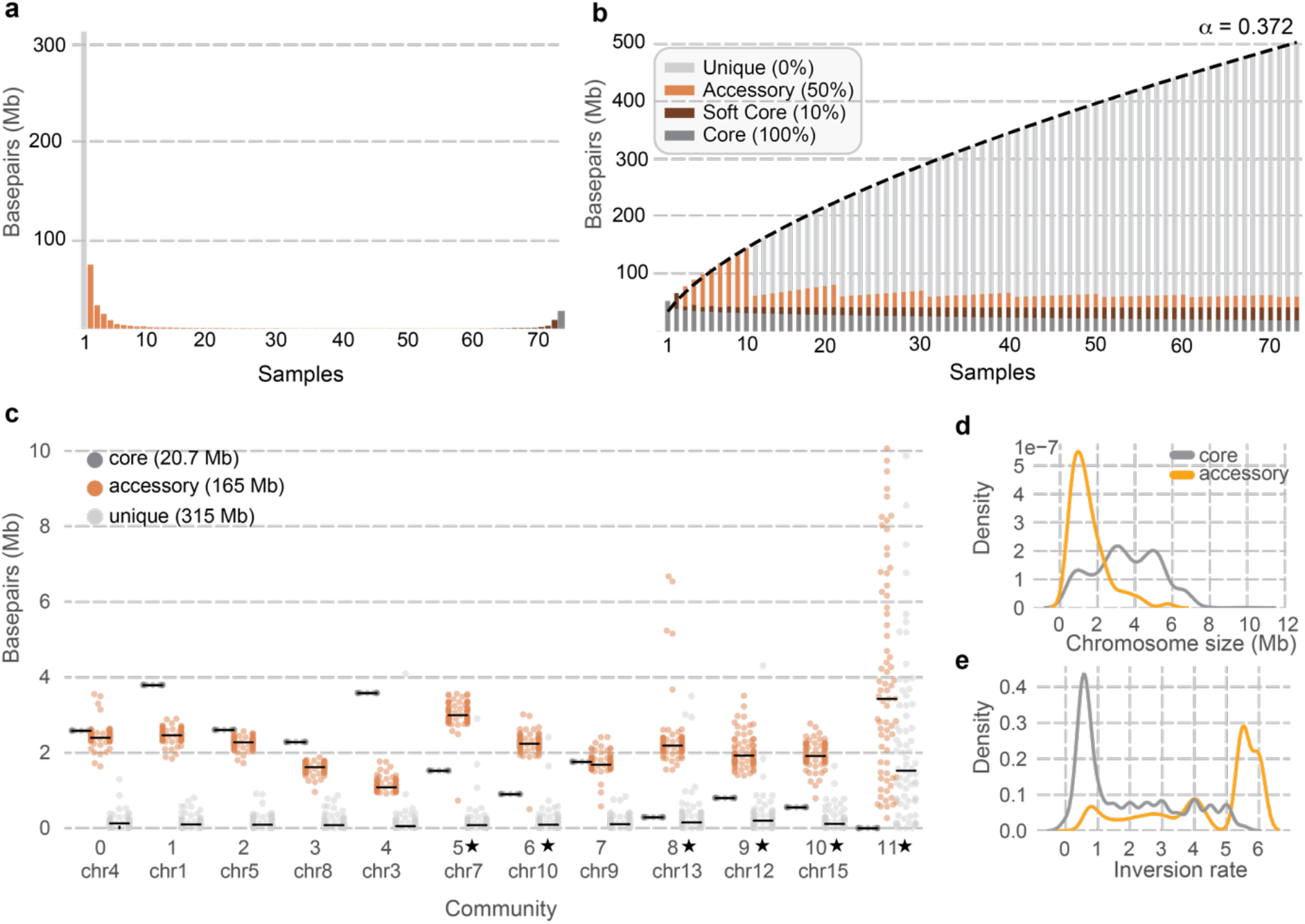
– Pangenome graph of *Fusarium oxysporum* is open and contains extensive accessory material. **a)** Histogram shows the amount of genetic material (in base pairs) that is present in an increasing number of samples (x-axis). Grey bar indicates the amount of material present in only one sample (unique material), the orange bars indicate the amount of material present in one to 65 strains, dark orange indicate the amount of material found in 65-72 strains, and dark grey shows the amount of material present in all strains (core). **b)** Pangenome growth graph of the complete *F. oxysporum* pangenome, as determined by Panacus (Parmigiani *et al*., 2024). Heaps Alpha < 1 indicates an open pangenome. **c)** Proportion (in base pairs) of core, accessory, and unique genetic material found in the different communities. Each dot represents a 500 bp window in the pangenome graph. As expected, community 11, containing all accessory chromosomes, lacks core material. Stars indicate the communities with more accessory than core material. **d)** Size of chromosomes in accessory community 11 (orange) compared to the size of chromosomes in the other core communities (grey). **e)** Inversion rate per 500 bp window in the pangenome graph in the core communities (grey) and accessory community 11 (orange). The inversion rate is determined by the number of inversions per genome and per node in the pangenome graphs.

To further investigate the expected co-linearity and variation between chromosomes in the pangenome graph, we analyzed the orientation and location of the nodes. We observed a higher number of inversions and nodes occur in varying order in the pangenome graph of the accessory community (Figure 2e, Fig S1). This shows that, while core chromosomes are largely co-linear and conserved, the sequence order between homologous regions in accessory chromosomes is not conserved.

To determine the amount of similarity between different strains, we determined the node-similarity in the pangenome graph. We observed that based on the core communities, most genomes are highly similar (average sharedness 80%). In the accessory community the average sharedness between genomes is lower (11%), in line with the observed variation and lack of core nodes. The similarity of the accessory regions between strains infecting the same host is higher for some formae speciales, such as the tomato infecting strains (median similarity of 46%, across 64 strains) (Fig S2), but not for all host specific strains. For example, we only observed 14% similarity across 71 banana infecting strains (Fig. S1). The high similarity in tomato infecting strains supports earlier reports showing that *F. oxysporum* strains infecting tomato contain a host-specific accessory chromosome that is associated with pathogenicity (Ma *et al*., 2010). The lack of similarity between accessory chromosomes in strains infecting other hosts suggests these might not have host-specific pathogenicity chromosomes, as we have recently proposed for banana-infecting strains (van Westerhoven *et al*., 2024a; Dijkstra *et al*., 2024).

### Accessory chromosomes can be grouped into separate co-linear communities

To further delineate subgroups of accessory chromosomes that share more extensive co-linearity, we performed an additional analysis where we only considered mappings between chromosomes that cover most of the corresponding chromosome, i.e., we exclude partial mappings between chromosomes. As a result, chromosomes are now grouped into 36 communities, composed of the same eleven core communities as well as 25 additional accessory communities, demonstrating that accessory chromosomes can be further separated into groups based on co-linearity between chromosomes. The retrieval of the same eleven core chromosome communities, even when excluding splitting, further supports that core chromosomes are colinear. Nevertheless, we observed 20 cases where a chromosome maps to two different communities. To construct the final pangenome graph, these chromosomes were manually split and added into the corresponding homologous community. Not all chromosomes present in the 73 *F. oxysporum* strains have a homologous chromosome in another isolate, we find 107 unique chromosomes, with an average size of 1.1 Mb, without any mappings to a homologous chromosome, suggesting that these are strains specific.

The separation of accessory chromosomes into smaller co-linear communities enabled us to limit our analysis to highly similar accessory chromosomes. Interestingly, we observed that some of the co-linear accessory chromosomes are unique to, or at least enriched for one of the formae speciales (Fig. 3c). For example, chromosome 14 of *Fol*4287 is well known to play a role in pathogenicity towards tomato (Ma *et al*., 2010), and has been found to be present in genetically diverse tomato infecting *F. oxysporum* strains (Fokkens *et al*., 2018). Similarly, we observed that community 13, containing *Fol4*287 chromosome 14, solely consists of tomato infecting *F. oxysporum* strains. The other five accessory chromosomes of *Fol*4287 (chromosomes 2, 6, 11, 16, and 17) are found in accessory community 12. The grouping of these five accessory chromosomes supports previous reports that suggested that the accessory chromosomes in *F. oxysporum* strain *Fol*4287 are highly similar, due to extensive segmental duplications (Ma *et al*., 2010; van Westerhoven *et al*., 2024a).

**Figure 3.**
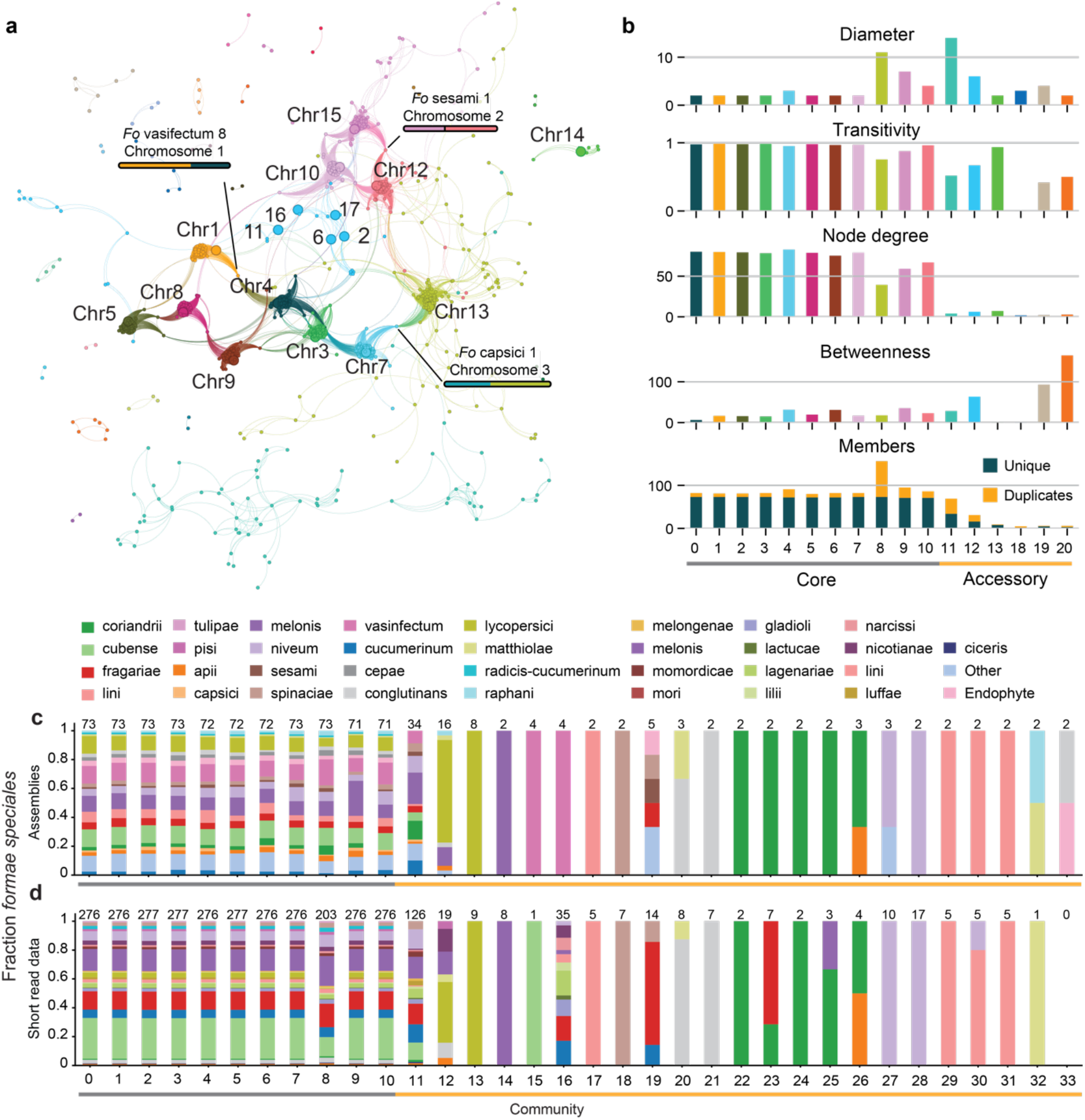
– Accessory chromosomes can be separated into several co-linear communities, revealing host-specific accessory chromosomes. **a)** The chromosome network displays all pair-wise mappings spanning the full length of chromosomes. Nodes represent chromosomes and edges represent mappings between chromosomes. Colors indicate the different communities that are detected in the network. Chromosome names correspond to the chromosomes found in the *Fol*4287 reference genome assembly, and these chromosomes are depicted by larger nodes in the graph. Orange circles highlight accessory chromosomes. **b)** Network statistics of the different communities in the chromosome network. Colors correspond to the communities in the network, see (a). **c)** Distribution of the different strains and their host specificity over the different communities. All strains are found in the core communities (grey line, x-axis). The accessory communities (orange line, x-axis), on the other hand, contain a subset of strains and fifteen accessory communities are host specific; the number on top indicate the total number of genomes in the community. **d)** Distribution of different *F. oxysporum* strains and their host specificity over the different communities.

In total, we observed 15 clusters that are specific to a formae speciales (Fig. 3c), suggesting that these might also represent pathogenicity chromosomes. However, most formae speciales were only represented by a limited number of genome assemblies (Fig. 1b,c). To include a broader set of strains, we sought to detect presence of accessory communities in a set of genome assemblies based on short-read sequencing data. We used a collection of 588 genome sequences that had been assembled from public data available at the Sequence Read Archive (Table S3). To rapidly determine the chromosomes, present in one of these genome assemblies, we compared the k-mer profiles of these genome assemblies against the k-mer profiles of the chromosome communities. As to be expected, core communities are present in all strains and the accessory communities show a variable presence/absence pattern (Fig. 3d). Using the formae speciales known for 276 strains, we could distinguish 14 host specific chromosomes (present in more than four strains; Fig 3d), including the tomato-specific community 13 that contains *Fol*4287 chromosome 14. These analyses show that some accessory communities are indeed host specific in a wide variety of *F. oxysporum* strains, yet many other accessory communities are not restricted to a single host.

### Accessory chromosomes are a mosaic of accessory regions

To get a more detailed insight into the variation within homologous chromosomes, we constructed a pangenome variation graph per community. This resulted in a combined pangenome, consisting of 443 Mb of genetic material that is separated over 31,289,751 nodes and connected by 44,436,449 edges. This pangenome graph consists of 18.8 Mb core, 14 Mb soft-core (>90% of strains), 161 Mb accessory, and 250 Mb of unique material, and is open (Heaps Alpha = 0.41, Fig S3). As to be expected, the fraction of core material is considerably larger than in the previous pangenome graph (Fig. 2), as accessory communities now consist of more similar material (Fig. S3).

We observed that the accessory chromosomes contain a high number of inversions (Fig 2c) and are generally not co-linear (Fig. 3). We distinguished three different types of accessory communities. Most accessory communities (22) are small with only five chromosomes. The chromosomes in these clusters are overall well connected (betweenness: 30.5, diameter: 1.15) and share a large amount of genetic material (49 Mb core, 21 Mb accessory, and 27 Mb unique). In addition, we find two large accessory communities (community 11 and community 12 with 69 and 31 members, respectively). These accessory communities have a larger diameter (10.0) and a higher betweenness (45.9) than the small accessory communities, indicating that the nodes in the communities are loosely connected; not all chromosomes share genetic material, analogous to accessory community 11 in our previous analysis (Fig. 1). Moreover, the pangenome graph of community 11 and 12 does not contain any core nodes (Fig. S4), further supporting the lack of shared genetic material between the chromosomes in these communities.

To determine what links the chromosomes in these large and loosely connected accessory communities, we visualized the presence and absence of nodes in the pangenome graph (Fig. 4a). This shows that not all chromosomes share regions with each other but stretches of similar regions are found in different combinations (Fig. 4b). For example, different parts of an accessory chromosome (*Fol*029, chromosome 12) are present in other accessory chromosomes (Fig. 4c), indicating that accessory chromosomes share different accessory region. Together, these findings supports the idea that accessory chromosomes evolve through chromosomal rearrangements, resulting in a mosaic of genetic material from different sources (Fokkens *et al*., 2018).

**Figure 4.**
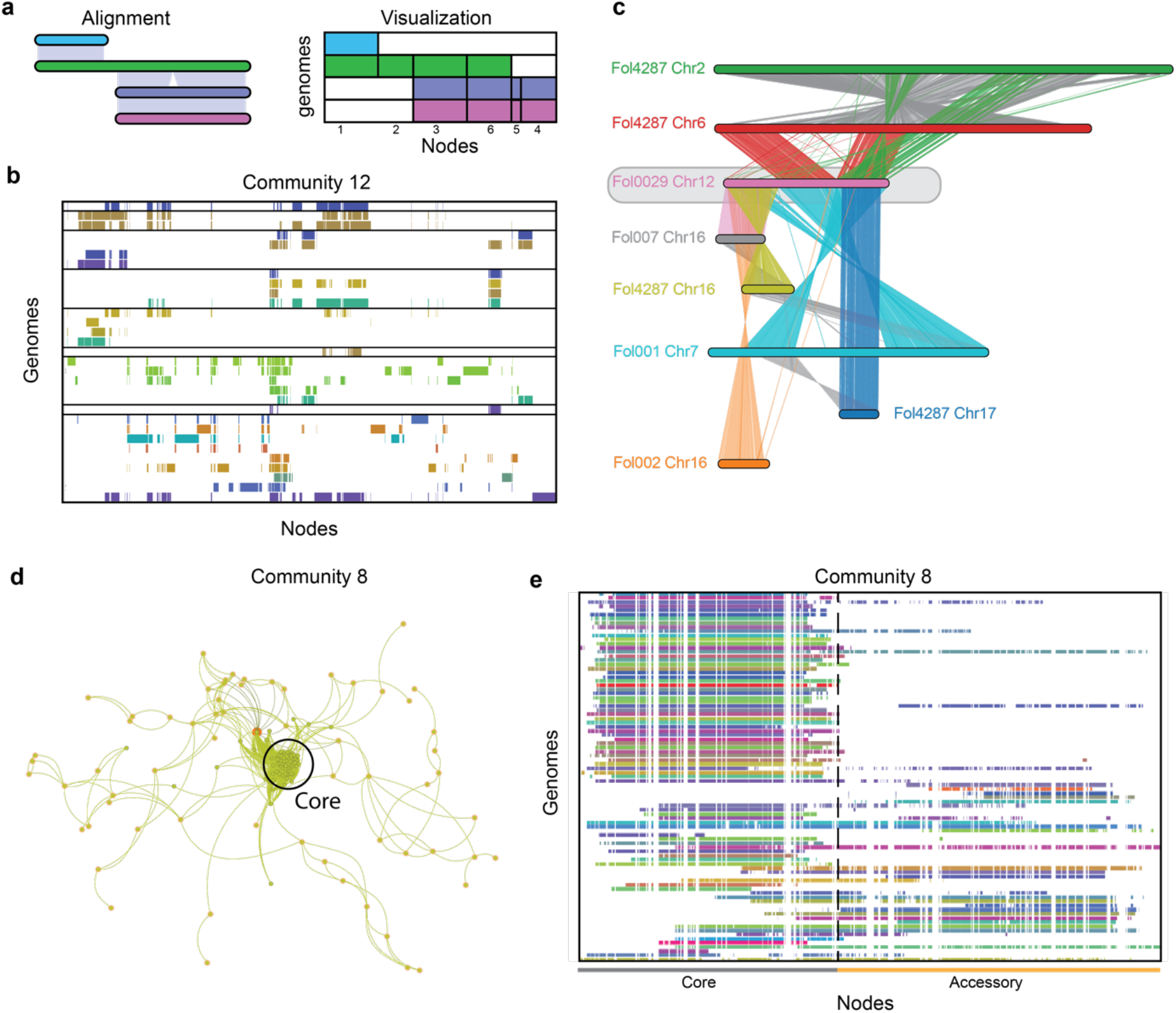
– Accessory chromosomes are a mosaic of accessory regions from different genomes. **a)** Translation from a pair-wise chromosome alignment to a pangenome graph presence/absence matrix. The visualization places the nodes in the pangenome graph on the x-axis and the genomes on the y-axis. Colored blocks indicate a node that is present in a genome, while white indicates a node that is absent in the respective genome. **b)** Presence/absence of nodes in the pangenome graph based on the 69 members of community 12. Note, none of the nodes are present in all chromosomes. **c)** Alignment of eight chromosomes found in community 12. All chromosomes are aligned to *Fol*0029 chromosome 12 (pink) and share different but similar genomic regions with the other chromosomes. The connecting lines indicated regions of shared similarity, colored according to query chromosome. Grey lines indicate similarity between two neighboring chromosomes, chromosome 2 and chromosome 6 of *Fol*4287 are homologous. **d)** Community 8 shows a collection of core chromosomes surrounded by various accessory chromosomes (highlighted in orange). **e)** Node presence/absence matrix of a subset of chromosomes in community 8. The node presence and absence clearly demonstrate that the core part of the chromosomes is similar across all 73 genomes (grey line). Importantly, some core chromosomes contain an accessory region, while in other cases the accessory regions are separate from the core chromosome.

Next to these two types of accessory communities, we also identified accessory chromosomes present in two core chromosome communities (community 8 and community 9 with 63 accessory and 14 accessory chromosomes, respectively; Fig. 4d). This pattern can indicate that accessory chromosomes originated from core chromosomes or that accessory chromosomes fused to core chromosomes. We observed that similar core chromosomes are present in community 8, as well as additional separate accessory chromosomes that do not share homology to the core regions (Fig. 4d,e). However, this accessory region is attached to the core chromosome in some strains, whilst in others the accessory region is separate (Fig. 4e). This demonstrates that the link between core and accessory chromosomes in this community is caused by the fusion of an accessory region to a core chromosome, an arrangement that has been previously observed in various *F. oxysporum* genome assemblies (Ma *et al*., 2010; van Westerhoven *et al*., 2024a; Dijkstra *et al*., 2024).

### Accessory chromosomes can be transferred between genetically different *F. oxysporum* strains

Accessory chromosomes can be transferred between *F. oxysporum* strains as has been experimentally shown for a few chromosomes in some strains (Ma *et al*., 2010; van Dam *et al*., 2017; Li *et al*., 2020) Horizontal transfer can also occur in natural populations and has been identified based on the presence of similar accessory chromosomes between distantly related strains (van Dam *et al*., 2016; Fokkens *et al*., 2018; Henry *et al*., 2021). To determine horizontal transfer events in our dataset, we analyzed the similarity of accessory chromosomes between strains from different taxonomic clades. We observed that ten communities contain members from different taxonomic clades of the *F. oxysporum* species complex (Fig. 5a), suggesting that horizontal transfer might have taken place. For instance, community 11 contains strains from the taxonomic clade 1, 2, and 3. One strain from clade 1 (*Foc*013) shares 58% similarity with clade 2 genome *Fom*0021, but only 12% similarity with closest clade 1 neighbor (*Fo*4, Fig. S5), indicating that horizontal transfer has occurred between strains from clade 1 and clade 2.

**Figure 5.**
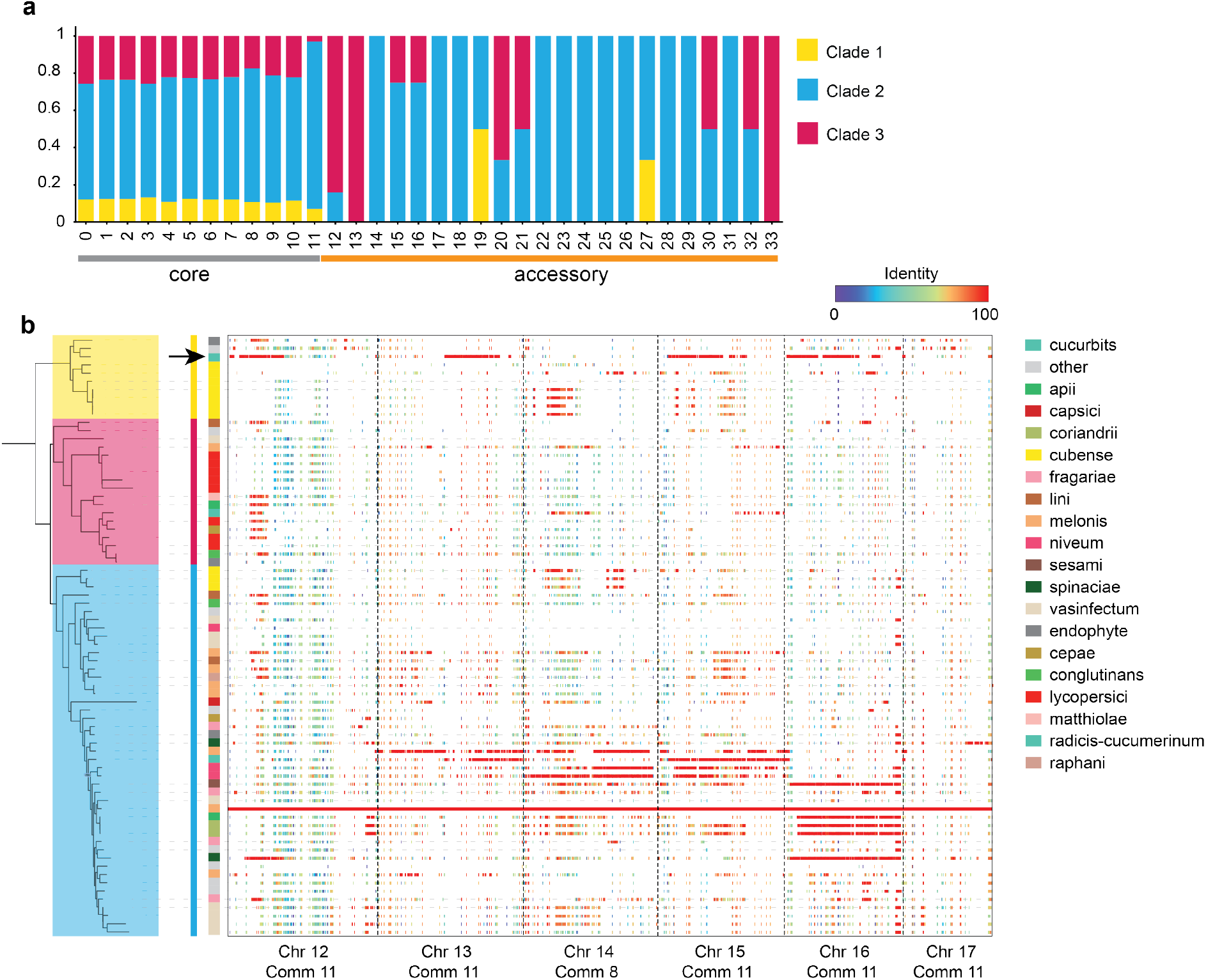
– Similar accessory chromosomes can be found in different *Fusarium oxysporum* clades. **a)** Distribution of *F. oxysporum* clades over the different chromosome communities (x-axis). Ten accessory communities contain strains from different taxonomic clades, indicating that similar chromosomes can be shared by *F. oxysporum* strains from with large phylogenetic distances, even from different taxonomic clades. **b**) Similarity of six accessory chromosomes found in the melon infecting *F. oxysporum* strain *Fom*0021 (x-axis) across all 73 *F. oxysporum* strains (y-axis), ordered based on the phylogenic relationship of the *F. oxysporum* strains (left). Only alignments that span at least 5kb are shown. The color indicates the percent identity. Accessory regions from *Fom*0021 are found in different strains in clade 2 as well as in one strain (*Foc*013) in clade 1, highlighted by an arrow.

To get a more detailed insight into the traces of horizontal chromosome transfer between *Fom*0021 and *Foc*013, we performed whole-genome alignments between the accessory chromosomes found in *Fom*021 to all 73 genomes. Interestingly, the accessory chromosomes are present in some but not all of the most closely related species, and similar genomic regions could even be found in a strain from different taxonomic clades. For example, chromosome 12, 13, 14 and 16 from *Fom*0021 (formae speciales *melonis*) are also found in *Foc*013 (formae speciales *cucumerinum*) from clade 1 and a region of accessory chromosome 14 of *Fom*0021 can be found in four banana infecting genomes (formae speciales *cubense*) in clade 1. This further corroborates that accessory chromosomes are shared across genomes and further highlights that horizontal transfer occurs between clades of *F. oxysporum*. Interestingly, the shared genomic regions are present in strains infecting different hosts, suggesting that the transfer is not always associated with the same host range.

## Discussion

Pangenome variation graphs offer a reference-free genome representation of different individuals of the same species. This can provide insights into the genome organization, and can improve downstream genome-based analysis such as variant calling, and gene-annotation (Li *et al*., 2022; Wang *et al*., 2022; Guarracino *et al*., 2023; Skiadas *et al*., 2024). Such reference free analysis would be especially useful for analyzing accessory chromosomes in filamentous plant pathogens, as these are not necessarily present in the reference genome assembly. Moreover, accessory chromosomes are often highly diverse hampering reference-based analyses (Sherman & Salzberg, 2020; Barragan *et al*., 2024). By combining a homologous chromosome mapping strategy with the construction of a pangenome variation graph, we captured all accessory chromosomes in a collection of 73 *F. oxysporum* strains and obtained insights into their evolutionary dynamics. We found eleven conserved core chromosomes and a large number of different accessory chromosomes, which evolve through rearrangements that result in a mosaic of different accessory genomic regions. Moreover, we found evidence for horizontal transfer of accessory chromosomes between naturally occurring *F. oxysporum* strains from different phylogenetic clades. These results highlight that pangenome variation graphs can be successfully reconstructed for species carrying accessory chromosomes and can guide the exploration of genetic variation that underlies specific phenotypes and provides insights into their genome organization to obtain insights into genome evolution.

*F. oxysporum* is a highly diverse species complex that infects a wide variety of hosts (Edel-Hermann & Lecomte, 2019; Fayyaz *et al*., 2023; Armer *et al*., 2024). We identified an open pangenome, meaning that every new strain adds yet unseen additional genomic regions to the pangenome, which is in line with previous gene-based pangenomes in *F. oxysporum* (Fayyaz *et al*., 2023; van Westerhoven *et al*., 2024a) and is expected for a highly diverse species (Tettelin *et al*., 2005). The rapid diversification of accessory chromosomes can underly the observed diversity (Dong *et al*., 2015; Yang *et al*., 2020). Using the pangenome variation graph, we identified a large set of diverse accessory chromosomes with a mosaic composition, encoding a unique combination of accessory regions, similar to what has been observed in the accessory genome of other fungi (Fokkens *et al*., 2018; Langner *et al*., 2021; Barragan *et al*., 2024). Moreover, similar accessory chromosomes are found between strains from different clades, highlighting that accessory chromosomes can be horizontally transferred between distantly related strains. This supports previous studies suggesting that clades are not genetically isolated in *F. oxysporum* (van Westerhoven *et al*., 2024b), and that genetic material can be exchanged not only within but also between taxonomic clades. The horizontal transfer of accessory chromosomes, together with rearrangement of accessory material, can provide genetic diversification that is especially important in asexual fungal species (Seidl & Thomma, 2014; Fokkens & Rep, 2023).

In *F. oxysporum*, many accessory chromosomes carry important pathogenicity genes and play a role in determining the host range (Ma *et al*., 2010; van Dam *et al*., 2017; Armitage *et al*., 2018). Similar to previous studies, we found highly similar and host-specific accessory chromosomes within several formae speciales. However, host-specific accessory chromosomes are not identified for all formae speciales. It remains unclear why some formae speciales have a clear pathogenicity related accessory chromosomes, like tomato (Fokkens *et al*., 2018), whereas others have variable accessory chromosomes (Henry *et al*., 2021; van Westerhoven *et al*., 2024a). This variability suggests that some host-specificities arose multiple times independently, or the genes required for this host specificity are reshuffled between accessory chromosomes or between accessory chromosomes and the core. Additionally, limited phenotyping data makes it difficult to study host specificity. In general, formae speciales are assigned based on the isolation source, but a strain may infect multiple hosts or cause different symptoms in the same host (Henry *et al*., 2021; Batson *et al*., 2021). The pangenome constructed in this study can serve as a valuable resource for exploring chromosome diversity within and between *formae speciales* of *F. oxysporum*. This can enable the identification of conserved regions and essential pathogenicity genes, providing insights into the host range of strains. Additionally, the pangenome can place newly sequenced and assembled *F. oxysporum* strains into a larger context, this can reveal the presence of accessory chromosomes, show what accessory regions are shared and thereby provide an important framework to clarify the role of accessory regions in pathogenicity.

Different forms of pangenomes can be constructed, based on genes, whole-genomes, or k-mer content (Andreace *et al*., 2023). The optimal approach depends on the biology of a species, as well as the exact biological question to be addressed. Research on pangenomes in fungi thus far mainly focused on gene-based pangenomes, representing all genes within a species (Badet *et al*., 2020; Barber *et al*., 2021). This provides important insights into the evolution of gene content, especially of pathogenicity genes, but typically offers limited information on the genome organization and the presence and shape of accessory chromosomes. Insights into accessory chromosome dynamics have been previously obtained by pairwise or reference-based comparisons (Feurtey *et al*., 2023; van Westerhoven *et al*., 2024a; Barragan *et al*., 2024). The here applied reference-free pangenome variation graph enables large-scale comparisons of accessory chromosomes and can be applied in different fungi to obtain insights into chromosome dynamics. Moreover, a pangenome variation graph is valuable data by improving variant calling (Sirén *et al*., 2021) as well as annotation of genes and transposable elements (Fiddes *et al*., 2018; Skiadas *et al*., 2024; Groza *et al*., 2024), thereby enhancing our capabilities to study a species’ biology. Future analyses of pangenome variation graph can thus not only help to reveal insights into the evolution of the genome organization, but also can help to reveal new insights into the population structure, pathogenicity genes, and other drivers of phenotypic variations.

## Material and Methods

### DNA isolation and whole-genome sequencing and assembly

For strains Fol005, Fol007, Fom011, Fom014, Fom021, Fom024 and Fom025, strains were taken from glycerol stock, grown on PDA for three days at 25°C, after which 100 ml of NO3 was inoculated with an overgrown agar plug. Samples were grown in this liquid culture for 5-7 days at 25°C and 150 RPM. Mycelium was harvested by filtering through miracloth, stored in liquid N2 and freeze-dried overnight. Per sample DNA was isolated from ∼250 mg of ground mycelium using multiple rounds of phenol-chloroform extractions, two rounds of chloroform extractions, and was precipitated twice. Quality and quantity of DNA was checked with Nanodrop, Qubit and agarose gels. DNA samples were sequenced at KeyGene on three PromethION FLO_PRO002 cells where basecalling was performed with the high-accuracy model in MinKNOW 4.2.6 and reads with quality q >= 7 were selected. Adapters were trimmed with poreChop, and reads were filtered based on their quality and length with Filtlong (version 0.2.0, --min_length 1000, --min_mean_q 80 en --min_window_q 70). Per strains, the 80% longest reads were selected and assembled with Flye (v2.8). Strains Fol029 and FolMN25 were sequenced and assembled according to <u>van Dam *et al*., 2017</u>.

### Genome selection

*Fusarium oxysporum* genome assemblies have been downloaded from NCBI (July 2024), retaining genome assemblies with less than a 100 contigs and an N50 larger than 2 Mb. In addition to the genomes from NCBI, we included twelve previously assembled whole-genome assemblies (Table S1). Not all genomes have been assembled and processed in the same way. To account for this heterogeneity, we applied a homogenous filtering strategy and excludes fragmented chromosomes. We filtered contigs smaller than 50 kb, removing between 0 and 80 contigs per genome assembly (Table S1). We then accessed the genome assembly quality using Quast version 5.0.2 (Gurevich *et al*., 2013) and determined the presence of telomeric repeats using tidk version 0.2.41 (Brown *et al*., 2023). Furthermore, we constructed a phylogenetic tree of the strains using conserved single-copy BUSCO genes with the BUSCO phylogenomic pipeline, available at https://github.com/jamiemcg/BUSCO_phylogenomics.

### Network construction

To detect groups of homologous chromosomes, we first mapped all contigs to each other using wfmash version 0.10.3 (Marco-Sola *et al*., 2021). To determine the optimal settings for *F. oxysporum*, we tried different parameters for the length of mapped segments (-s, 2.5kb, 5kb, and 10kb) and the block length the chained segments (-l, from 5kb, 10kb, 25kb,50kb, and 100kb); increasing segment and block length also increases the length of co-linear blocks between chromosomes. The resulting all-vs-all homology mappings between chromosomes were used to construct a chromosome network. In addition to this network, we also created a network with splitting of chromosomes disabled (wfmash flag -N), removing partial chromosome mappings. These two all-vs-all homology mappings were treated similar for downstream comparisons.

Chromosomes communities were detected in the all-vs-all homology mappings using the Leiden Algorithm (Traag *et al*., 2019) as described by (Guarracino *et al*., 2023; Garrison *et al*., 2024). The chromosome homology networks were visualized using Gephi version 0.10 (Bastian *et al*., 2009), and the network layout was calculated using the YiFan Hu algorithm. Community statistics were determined using iGraph python module version 0.10.8 (Csárdi & Nepusz, 2006). To understand the chromosome network in relation to the *F oxysporum* reference genome assembly *Fol*4287, we plotted the communities assigned to chromosomes in the *Fol*4287 using pafR version 0.0.2 (https://dwinter.github.io/pafr/).

### Splitting of chromosomes

Before we could construct the pangenome graphs per community, we needed to ensure that each community represent homologous chromosomes. However, in some cases one chromosome maps to two different communities, for example due to chromosome fusion or errors during genome assembly. To include chromosome that map to two different communities, we split these chromosomes at the breakpoint and assigned chromosomal regions to the corresponding communities. Chromosomes were split into two different communities when a consecutive stretch of 1 Mb mapped to a different community and when this part occurred at the flank of the chromosome, and thus we only split large co-linear fractions of the chromosome, preventing us to artificially splitting chromosomes into too small sections. Some windows in the chromosome did not map to any community. To maintain the chromosome structure, these unmapped regions were ignored when we determine the flanks of a chromosome and, these unmapped regions were assigned to the new community when they flank the split region.

### Pangenome variation graph construction

We constructed a pangenome variation graph per community detected in the all-vs-all chromosome network. The pangenome graph was constructed with the PanGenomeGraphBuilder version 0.3.0 (PGGB, Garrison et al., 2024) with the following parameters: a segment length of 2,500 bp (-s), a block length of 50,000 bp (-l), a percent identity threshold of 95% (-p), and 72 haplotypes (n-1). The generated pangenome variation graph was then sorted and visualized using the optimized dynamic genome/graph implementation version 0.8.6.0 (odgi, Guarracino et al., 2022). The general graph statistics were calculated with odgi stats (-S), the pairwise genome similarity was assessed with odgi similarity, and the percentages of core and accessory nodes were determined using odgi stats (-a). To assess structural variation, the mean inversion rate was calculated using odgi bin -w 50,000 bp, and node presence/absence patterns were visualized with odgi viz. Subgraphs for different communities were combined using odgi squeeze to form the final *F. oxysporum* pangenome variation graph. Finally, this joined *F. oxyporum* variation graph was used to evaluate the pangenome growth curve with Panancus (version 0.2.3, Parmigiani et al., 2024).

### Short read genome assemblies and matching to the pangenome graph

To include a larger variety of *F. oxysporum* strains, we compared the k-mer profile of 588 fragmented *F. oxysporum* assemblies (Table S3) to the k-mer profile of the co-linear accessory communities, based on our unsplit mapping approach. The strains were assembled from short read sequencing data downloaded from NCBI using spades version 3.13.0 (Bankevich *et al*., 2012). To compare the k-mer profiles, we used sourmash gather (--bp-threshold 0), version 4.8.9 (Pierce *et al*., 2019), and determined the presence of a chromosome community in the *F. oxysporum* strains based on the percentage of k-mer from the chromosome community that matched to the genome assemblies. The k-mer profile of a community is based on all chromosomes in the community, including similar chromosomes, and thus the short read k-mers will always match only a small percentage of the community. We considered that the maximal percentage of k-mers from a genome assembly to match a community to represent the optimal mapping. This maximum mapping percentage is used to normalize the matching percentage across all genome assemblies, and a community is considered present in a genome assembly when at least 20% of the maximum percentage is successfully mapped.

### Analysis of accessory communities

To visualize similarity between accessory chromosomes we used PyGenomeViz version 1.0 (https://moshi4.github.io/pyGenomeViz/), in addition to the pangenome based visualizations from odgi viz (Guarracino *et al*., 2022). To visualize similarities to the Fom0021 reference genome, we used nucmer –maxmatch to align all assemblies to the Fom021 assembly. We filtered the alignments to only retrieve alignments with length larger than 5 kb and a precent identity higher than 90% using custom Python scripts (https://github.com/LikeFokkens/FOSC_multi-speed-genome).

## Data availability

Genome sequencing data and assemblies used in this study can be found in Table S1 and S3. Custom code generated for this study can be found https://github.com/Anouk-vw/FOSC_pangenome and https://github.com/LikeFokkens/FOSC_multi-speed-genome

## Acknowledgements

A.C.W, and G.H.J.K were supported by a grant from the Bill and Melinda Gates Foundation to the International Institute of Tropical Agriculture, Agreement No - AG-5797. The funders had no role in study design, data collection and analysis, decision to publish, or preparation of the manuscript.

