## Supplementary Figures 1 - 5 for "Reference-free identification and pangenome analysis of accessory chromosomes in a major fungal plant pathogen"

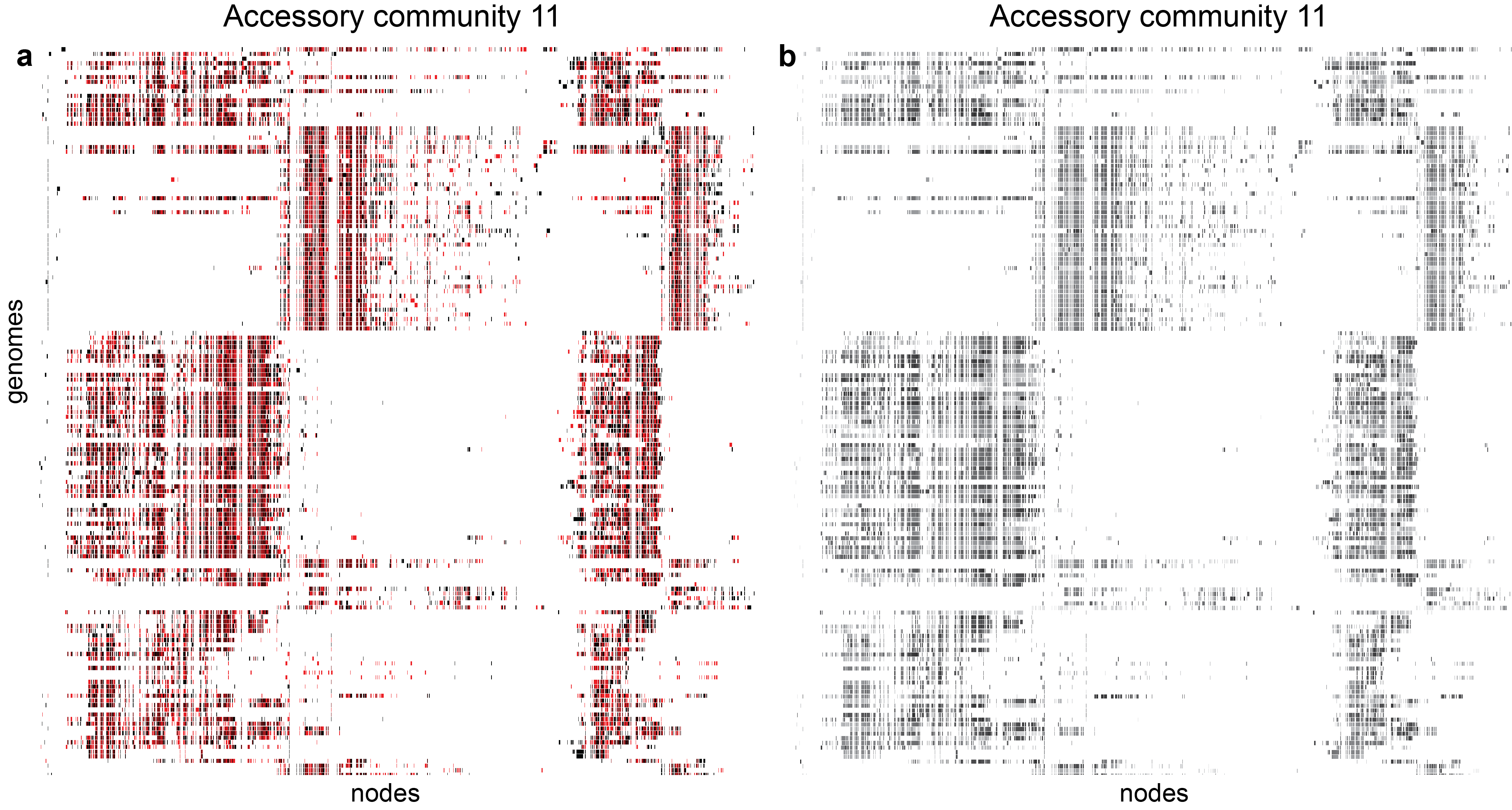

**Figure S1 – Inversions and path-orientation of community 11 (split) show lack of co-linearity.** Visualization of nodes that are present or absent in the pangenome of accessory community 11 (split). A subset of paths throught the graph are shown. Each row corresponds to a path (i.e., chromosome) in the pangenome graph showing nodes (sequences) as colored rectangles. matrix visualization created with odgi (Guarracino et al., 2022) **a**) Nodes are colored by the inversion rate per node. Red color indicates a node that is in reverse orientation in the path, black indicates the forward orientation of the node. Genomes contain many nodes in different orientation (forward and reverse), showing the high abundance of inversion in these chromosomes. **b)** Matrix visualization similar to (a), showing the location of a node in the linear chromosome. Light color indicates that the node is located at the beginning of the chromosome, dark color indicates that a note is located at the end of the chromosome. In co-linear chromosomes the nodes are oriented from dark to light. The mixed light and dark nodes of community 11 (split) suggest that these chromosomes are not co-linear.

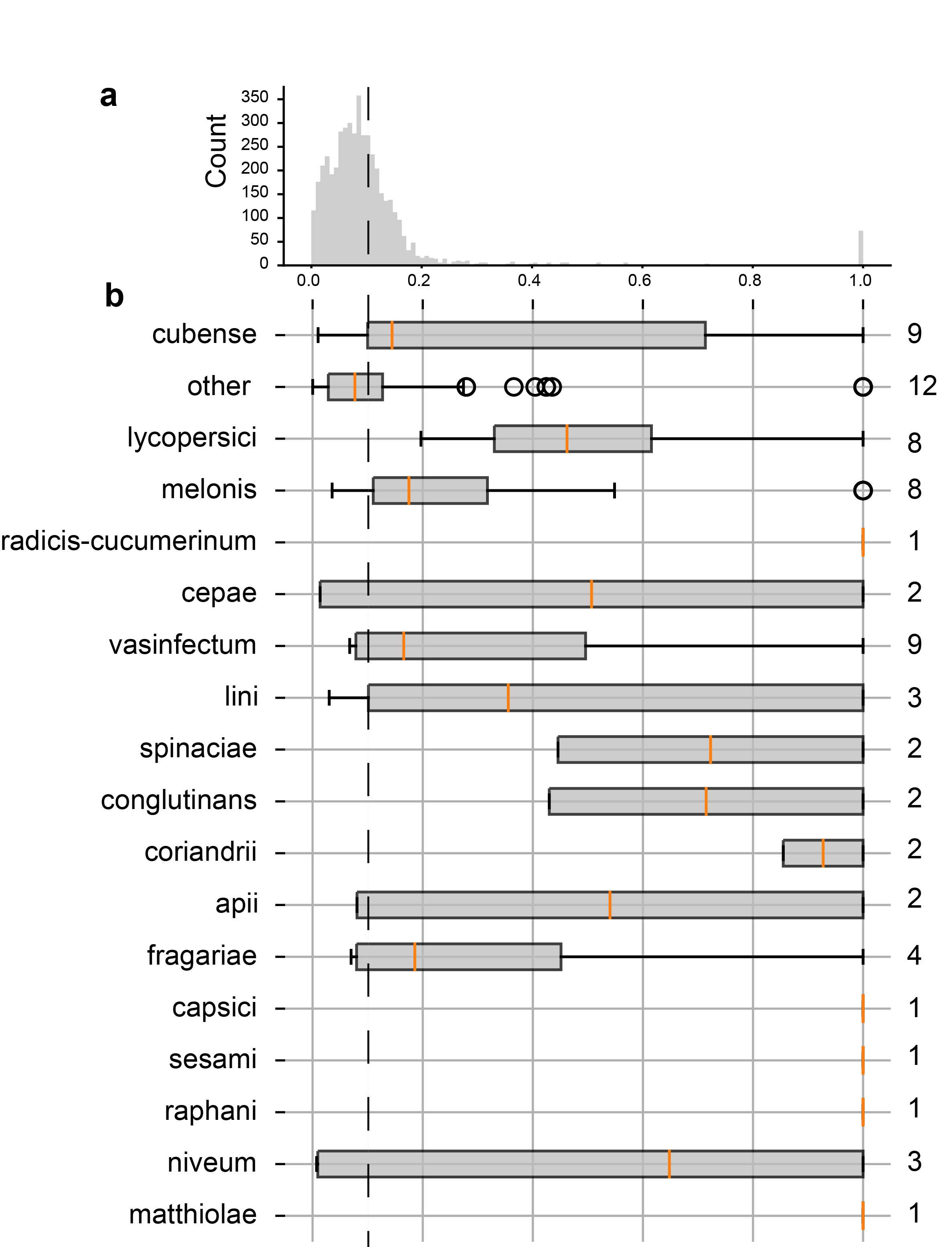

**Figure S2** – **Estimated identity between isolates infecting the same host in accessory community 11 (split).** **a)** Distribution of estimated pairwise identity across all genomes in community 11 (split). Community 11 (split) has a median pairwise identity of 16%. **b**) Estimated pairwise identity of isolates infecting the same host. Host specific isolates in general have a higher estimated pairwise identity then the median estimated identity across the entire community. The estimated pairwise identity between isolates infecting ‘other’ hosts is more similar to the distribution of pairwise identity across the community.

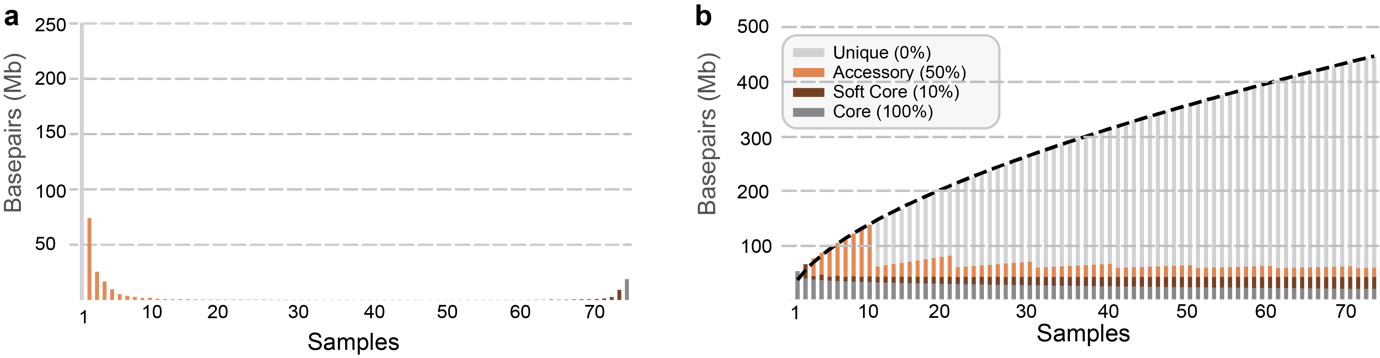

**Figure S3 –** **Statistics of the pangenome graph constructed based 36 homologous and co-linear chromosomes groups.** Pangenome graph is open and contains extensive accessory material a) Amount of genetic material (in base pairs) that is present in an increasing number of samples (x-axis). Grey bar indicates the amount of material present in only one sample (unique material), the orange bars indicate the amount of accessory material present in one to 65 strains, dark orange indicate the amount of soft-core material found in 65-72 strains, and dark grey shows the amount of core material present in all strains. b) Pangenome growth graph of the *F. oxysporum* pangenome, as determined using Panacus (Parmigiani et al., 2024). Heaps Alpha = 0.41, indicates an open pangenome.

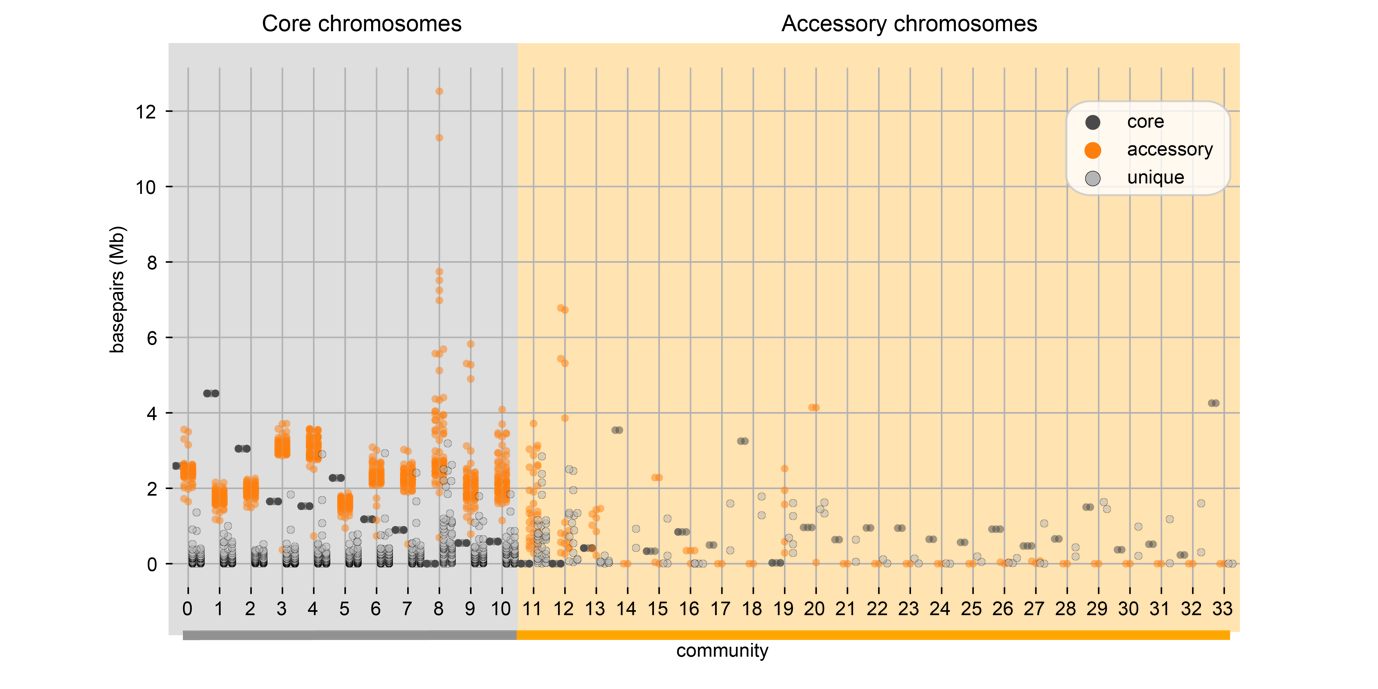

**Figure S4 - Proportion (in base pairs) of core, accessory, and unique genetic material found in the different communities.** Each dot represents a 500 bp window in the pangenome graph. First eleven communities are core chromosome communities (grey line) and contain core, accessory and unique material. Only community 8, containing 63 accessory chromosomes in addition to the 73 core chromosomes, does not encode any core material that is present in all chromosomes in this community. The accessory communities contain a variable amount of core material. The large accessory communities 11 and 12 do not contain any core material. The other accessory communities contain conserved genetic material that is present in all chromosomes in this community.

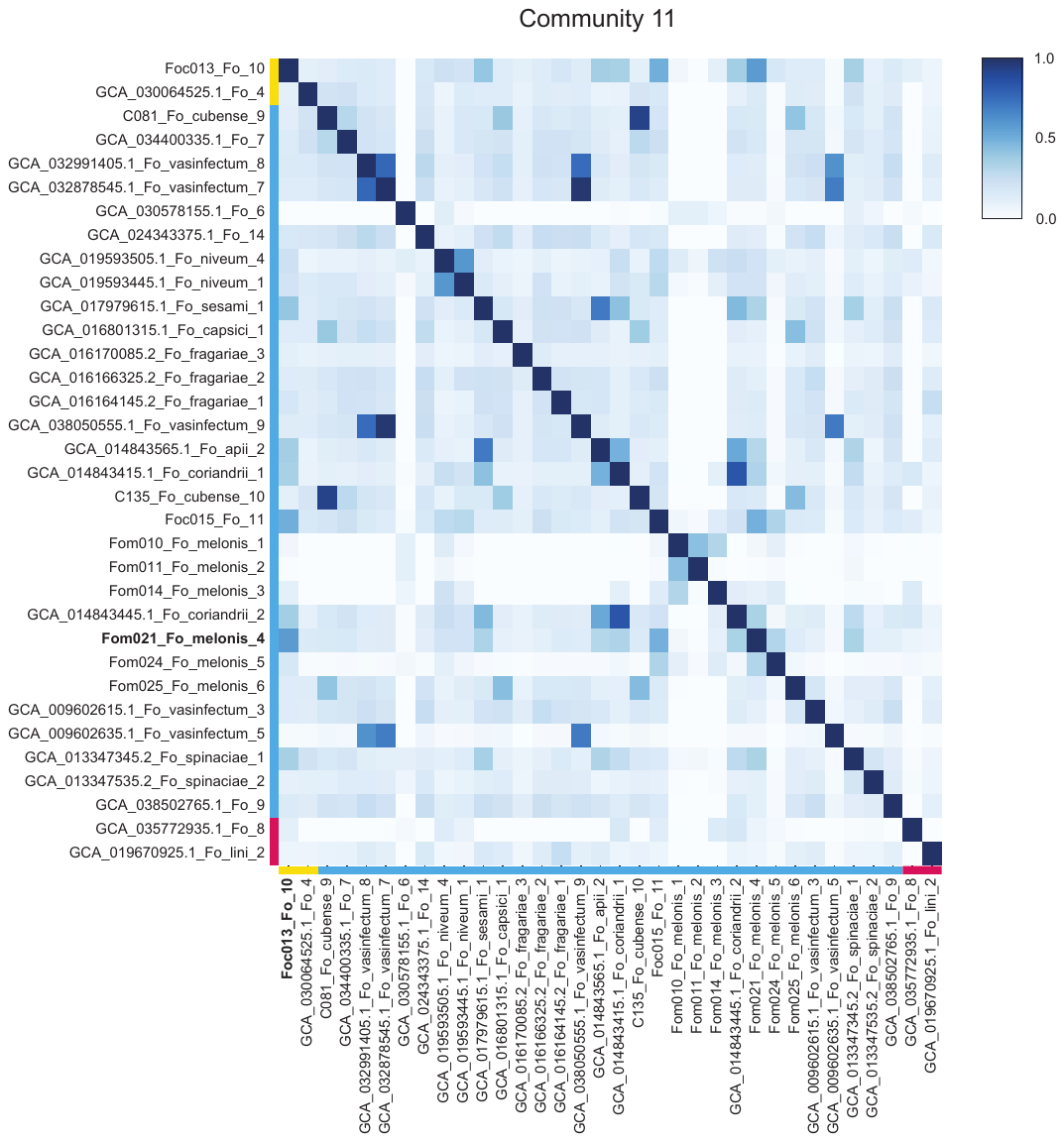

**Figure S5 – Estimated similarity of strains in community 11 (unsplit).** Color of the cell indicates pairwise similarity between isolates. Colored bars indicate the phylogenetic clade of the *Fusarium oxysporum* species complex to which this strain belongs to. The highest between clade similarity of 58% is observed between *Foc*013 (clade 1) and *Fom*0021 (clade 2), strain names highlighted in bold. Other strains, for example both banana infecting isolates (*cubense* 9 and *cubense* 10), and cotton infecting isolates (*vasifectum* 8, 9, and 10) have a remarkable high similarity.
